# Progressive Lineage Restriction of Bergmann Glia-like Progenitors during Postnatal Cerebellar Development

**DOI:** 10.64898/2026.06.09.731225

**Authors:** Toma Adachi, Kyoka Suyama, Sotaro Ito, Eriko Isogai, Masaki Sone, Mikio Hoshino

**Affiliations:** Department of Biochemistry and Cellular Biology, National Center of Neurology and Psychiatry (NCNP), Tokyo 187-8502, Japan; Graduate School of Medical and Dental Sciences, Science Tokyo, Tokyo 152-8550, Japan; Department of Biomolecular Science, Faculty of Science, Toho University, Chiba 274-8510, Japan

**Keywords:** Bergmann glia-like progenitors, postnatal cerebellum, lineage tracing, lineage restriction, molecular layer inhibitory neurons

## Abstract

Bergmann glia-like progenitors (BGLPs) are transient astroglial progenitors in the postnatal cerebellum, but how their lineage potential changes during development remains incompletely understood. Our previous electroporation-based study suggested that P0 BGLPs possess broader lineage potential than P6 BGLPs. Here, we performed recombination-based lineage tracing by cerebellar surface application of tamoxifen to *Ai9/+; Glast*^*CreERT2/+*^ mice and temporally analyzed the progeny of BGLPs labeled at P0, P3, P6, and P8. We found that BGLPs undergo progressive lineage restriction during postnatal development. P0 BGLPs gave rise to Bergmann glial cells (BGs), inner granule cell layer astrocytes (IGL astrocytes), white matter astrocytes (WM astrocytes), and molecular layer inhibitory neurons (ML-INs), confirming our previous electroporation-based findings. In contrast, P3 BGLPs generated BGs, IGL astrocytes, and WM astrocytes, whereas P6 BGLPs generated BGs and IGL astrocytes, and P8 BGLPs generated predominantly BGs. Thus, BGLP lineage output was progressively restricted from four progeny categories at P0 to a predominantly BG-restricted output by P8, suggesting that BGLPs dynamically adjust their cellular output during postnatal cerebellar maturation. Additional temporal analyses suggested that ML-INs are unlikely to be generated directly from P0 BGLPs, but may arise indirectly through astrocyte-like progenitors (AsLPs) and inhibitory neuron progenitors (INPs). These findings identify postnatal BGLPs as a useful *in vivo* model for studying progressive lineage restriction and stage-specific cellular supply during cerebellar development.

## Introduction

During development, progenitor cells often undergo progressive restriction of lineage potential, thereby generating distinct cell types in a stage-dependent manner (Koo et al., 2023; Yoon et al., 2018). Such temporal changes in progenitor competence are fundamental for producing the appropriate cellular composition of developing tissues (Koo et al., 2023; Yoon et al., 2018). In the nervous system, radial glial cells and other neural progenitors alter their lineage output as development proceeds, and this temporal regulation contributes to the orderly generation of neuronal and glial diversity (Telley et al., 2019; Koo et al., 2023). However, experimentally tractable *in vivo* systems in which a defined progenitor population undergoes stepwise lineage restriction within a short developmental window remain limited. Identifying such systems would provide an opportunity to understand how developmental time is linked to the progressive narrowing of progenitor potential.

The postnatal cerebellum provides a useful context for studying this issue because diverse neuronal and glial cell types are produced and positioned during a relatively short period after birth. In the cerebellar cortex, Bergmann glial cells (BGs), astrocytes in the inner granule cell layer (IGL astrocytes), astrocytes in the white matter (WM astrocytes), and molecular layer inhibitory neurons (ML-INs) are generated during postnatal development and contribute to the maturation of cerebellar structure and circuitry (Joyner and Bayin, 2022; Leto et al., 2006). Among the progenitor populations present during this period, Bergmann glia-like progenitors (BGLPs) are unique astroglial progenitors whose cell bodies are located in the Purkinje cell layer (PCL) and that extend unipolar processes toward the pial surface (Joyner and Bayin, 2022; Li et al., 2013). BGLPs proliferate mainly during early postnatal development and are no longer readily detected by approximately postnatal day 10 (P10) (Suyama et al., 2026). Thus, BGLPs represent a transient progenitor population whose lineage potential may be dynamically regulated during postnatal cerebellar maturation.

Previous studies have begun to define the lineage output of BGLPs. Recombination-based lineage tracing of P6 BGLPs showed that they generate BGs and IGL astrocytes, but not WM astrocytes or neuronal cell types (Cerrato et al., 2018). More recently, our electroporation-based labeling experiments suggested that P0 BGLPs have broader lineage potential than P6 BGLPs and can give rise to BGs, IGL astrocytes, WM astrocytes, and ML-INs (Suyama et al., 2026). These findings raised the possibility that BGLPs progressively lose lineage potential during postnatal development. However, because electroporation-based labeling is not a permanent recombination-based lineage-tracing method, the full lineage output of initially labeled BGLPs could not be rigorously defined. In addition, whether BGLP lineage potential changes in a stepwise manner across postnatal stages, rather than simply differing between P0 and P6, remained unknown.

In this study, we performed a temporal analysis of BGLP lineage potential using recombination-based lineage tracing by cerebellar surface application of tamoxifen to *Ai9/+; Glast*^*CreERT2/*^*+* mice. By labeling BGLPs at P0, P3, P6, and P8 and analyzing their progeny at later stages, we found that BGLPs undergo progressive lineage restriction during postnatal cerebellar development. P0 BGLPs generated BGs, IGL astrocytes, WM astrocytes, and ML-INs; P3 BGLPs generated BGs, IGL astrocytes, and WM astrocytes; P6 BGLPs generated BGs and IGL astrocytes; and P8 BGLPs generated predominantly BGs. Thus, the number of progeny categories generated by BGLPs decreased stepwise as development proceeded. Additional temporal analyses suggested that ML-IN production from P0 BGLPs is unlikely to occur directly, but may proceed indirectly through astrocyte-like progenitors (AsLPs) and inhibitory neuron progenitors (INPs). These findings identify postnatal BGLPs as an *in vivo* model for studying progressive lineage restriction during cerebellar development.

## Results

### Cerebellar surface tamoxifen labeling predominantly marks BG-like cells in early postnatal *Ai9/+; Glast*^*CreERT2/+*^ mice

To establish cerebellar surface Tx application as a tool for temporal lineage analysis, we first examined which cell types were initially labeled at early postnatal stages. BGLPs are astroglial progenitors that persist in the mouse cerebellum until approximately P10 (Suyama et al., 2026). Our previous electroporation-based analyses suggested that P6 BGLPs generate BGs and IGL astrocytes, whereas P0 BGLPs generate not only BGs and IGL astrocytes but also WM astrocytes and ML-INs (Suyama et al., 2026). However, because this method was not based on Cre-mediated recombination, it could not reliably capture all descendants of initially labeled cells. Cerrato and colleagues previously showed that application of tamoxifen crystals (tamoxifen citrate, Tx) to the cerebellar surface of P6 *Glast*^*CreERT2/+*^; *R26R*^*Confetti/+*^ mice (Mori et al., 2006; Betizeau et al., 2013) predominantly labels BG-like cells (BGLPs and BGs) (Cerrato et al., 2018). We therefore crossed *Glast*^*CreERT2/+*^ mice (Mori et al., 2006) with *Ai9* reporter mice (Madisen et al., 2010) and applied Tx to the cerebellar surface of P6 or P0 pups to determine the cell types generated from BGLPs in a recombination-based manner (Figures 1 and 2).

**Figure 1.**
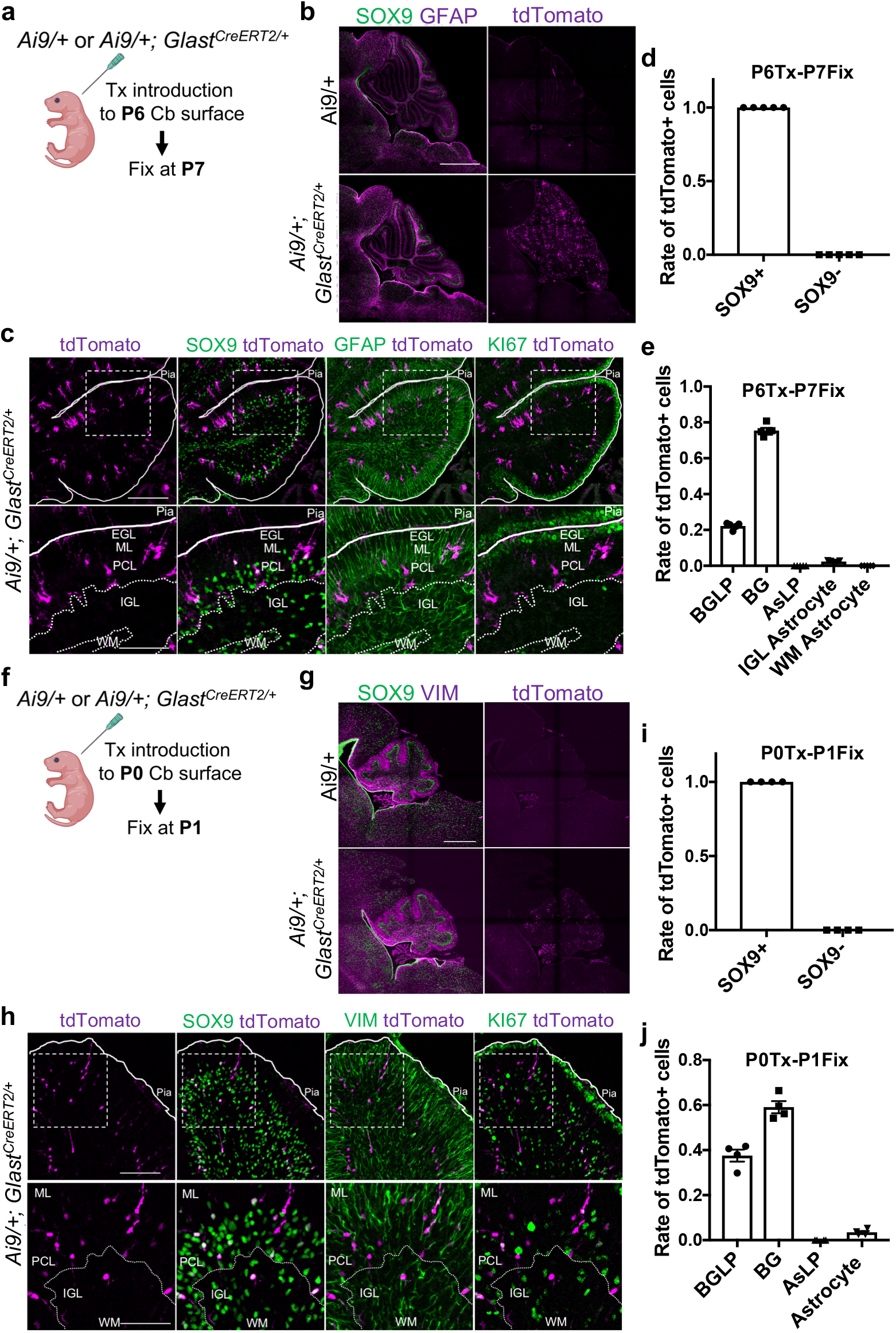
Cerebellar surface tamoxifen labeling predominantly marks BG-like cells in early postnatal *Ai9/+; Glast*^*CreERT2/+*^ mice. (a) Experimental scheme. Tamoxifen crystals (Tx) were applied to the cerebellar surface of *Ai9/+* or *Ai9/+; Glast*^*CreERT2/+*^ mice at P6, and the mice were fixed at P7. (b, c) Immunostaining of cerebella from tamoxifen-treated *Ai9/+ and Ai9/+; Glast*^*CreERT2/+*^ mice for DAPI, SOX9, GFAP, and KI67. Low-magnification images are shown in (b), and high-magnification images are shown in (c). tdTomato-positive cells were detected exclusively in the Purkinje cell layer (PCL) and molecular layer (ML) of *Ai9/+; Glast*^*CreERT2/+*^ mice. N = 5 mice per genotype. Scale bars: 400 μm in (b), 100 μm for the upper panels of (c), and 50 μm for the lower panels of (c). (d) Quantification of SOX9-positive cells among tdTomato-positive cells. N = 5 mice. (e) Quantification of the identities of tdTomato-positive cells. BGLPs were defined as SOX9+GFAP+KI67+ cells with unipolar processes located in the PCL and ML. BGs were defined as SOX9+GFAP+KI67− cells with unipolar processes located in the PCL and ML. AsLPs were defined as SOX9+GFAP+KI67+ cells with multipolar processes located in the WM. IGL astrocytes were defined as SOX9+GFAP+KI67− cells with multipolar processes located in the IGL. WM astrocytes were defined as SOX9+GFAP+KI67− cells with multipolar processes located in the WM. N = 5 mice. (f) Experimental scheme. Tx were applied to the cerebellar surface of *Ai9/+* or *Ai9/+; Glast*^*CreERT2/+*^ mice at P0, and the mice were fixed at P1. (g, h) Immunostaining of cerebella from tamoxifen-treated *Ai9/+ and Ai9/+; Glast*^*CreERT2/+*^ mice for DAPI, SOX9, VIMENTIN (VIM), and KI67. Low-magnification images are shown in (g), and high-magnification images are shown in (h). tdTomato-positive cells were detected exclusively in the PCL and ML of *Ai9/+; Glast*^*CreERT2/+*^ mice. N = 4 mice per genotype. Scale bars: 300 μm in (g), 100 μm for the upper panels of (h), and 50 μm for the lower panels of (h). (i) Quantification of SOX9-positive cells among tdTomato-positive cells. N = 4 mice. (j) Quantification of the identities of tdTomato-positive cells. BGLPs were defined as SOX9+VIM+KI67+ cells with unipolar processes located in the PCL and ML. BGs were defined as SOX9+VIM+KI67− cells with unipolar processes located in the PCL and ML. AsLPs were defined as SOX9+VIM+KI67+ cells with multipolar processes located outside the PCL and ML. Astrocytes were defined as SOX9+VIM+KI67− cells with multipolar processes located outside the PCL and ML. N = 4 mice. All data are shown as mean ± SEM.

**Figure 2.**
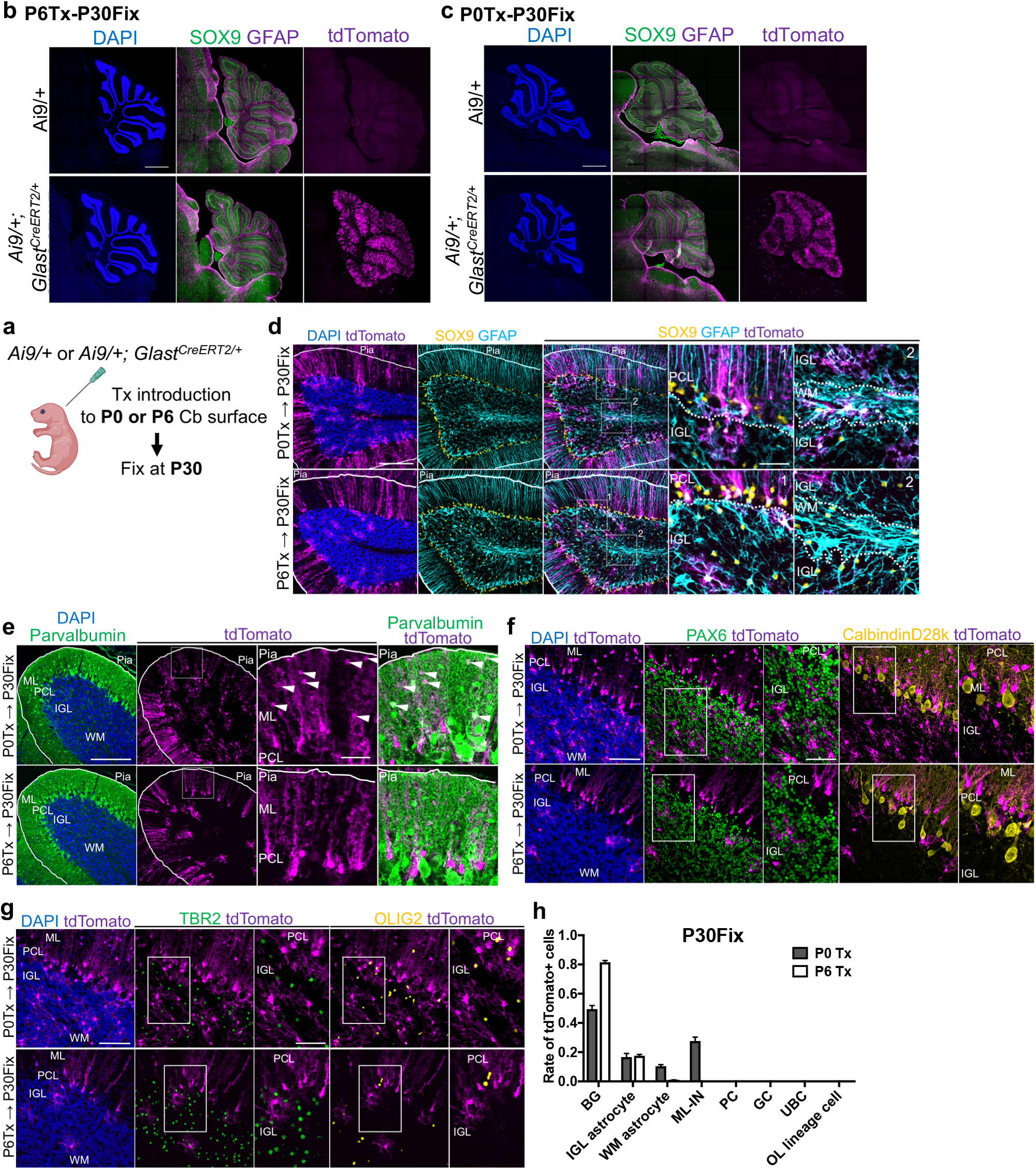
Recombination-based lineage tracing confirms broader lineage output of P0 than P6 BGLPs. (a) Experimental scheme. Tx were applied to the cerebellar surface of *Ai9/+* or *Ai9/+; Glast*^*CreERT2/+*^ mice at P0 or P6, and the mice were fixed at P30. (b–d) Immunostaining of cerebella from tamoxifen-treated *Ai9/+ and Ai9/+; Glast*^*CreERT2/+*^ mice for DAPI, SOX9, and GFAP. Low-magnification images are shown in (b, c), and high-magnification images are shown in (d). tdTomato-positive cells were detected exclusively in the *Ai9/+; Glast*^*CreERT2/+*^ mice treated with Tx at either P6 (b) or P0 (c). N=6 mice per genotype. Scale bars: 600 μm in (b, c), 100 μm for the left panels of (d), and 30 μm for the right panels of (d). (e–g) Immunostaining of cerebella from tamoxifen-treated *Ai9/+; Glast*^CreERT2/+^ mice for DAPI and Parvalbumin (e), DAPI, PAX6, and CalbindinD28k (f), or DAPI, TBR2, and OLIG2 (g), together with tdTomato. These analyses assess whether tdTomato-positive cells adopt molecular layer inhibitory neuron (ML-IN) (e), granule cell (GC) or Purkinje cell (PC) identities (f), or unipolar brush cell (UBC) or oligodendrocyte lineage cell (OL-lineage cell) identities (g). Scale bars: 100 μm for the left panels of (e), 30 μm for the right panels of (e), 50 μm for the left panels of (f, g), and 30 μm for the right panels of (f, g). (h) Quantification of the identities of tdTomato-positive cells. BGs were defined as SOX9+GFAP+ cells with unipolar processes located in the PCL and ML. IGL astrocytes were defined as SOX9+GFAP+ cells with multipolar processes located in the IGL. WM astrocytes were defined as SOX9+GFAP+ cells with multipolar processes located in the WM. ML-INs were defined as Parvalbumin+ cells located in the ML. PCs, GCs, UBCs, and OL-lineage cells were defined as CalbindinD28k+, PAX6+, TBR2+, and OLIG2+ cells, respectively. N=6 mice. All data are shown as mean ± SEM.

We first examined which cell types were labeled by this method at each developmental stage (Figure 1). When Tx was applied to the cerebellar surface of P6 *Ai9/+* or *Ai9/+; Glast*^*CreERT2/+*^ mice and the brains were analyzed 1 day later (Figure 1a), tdTomato-positive cells were detected exclusively in *Ai9/+; Glast*^*CreERT2/+*^ mice, but not in control *Ai9/+* mice (Figure 1b). All tdTomato-positive cells were SOX9-positive astroglial lineage cells (Figure 1c, d). Most tdTomato-positive cells were BG-like cells (BGLPs and BGs; SOX9+GFAP+ unipolar cells located in the PCL and ML; 97.7 ± 0.6%) (Figure 1c, e), whereas a small fraction were IGL astrocytes (SOX9+GFAP+KI67− multipolar cells located in the IGL; 2.3 ± 0.2%) (Figure 1c, e). Notably, AsLPs (SOX9+GFAP+KI67+ multipolar cells) were not labeled (Figure 1c, e).

We next performed the same analysis at P0. When Tx was applied to the cerebellar surface of P0 *Ai9/+* or *Ai9/+; Glast*^*CreERT2/+*^ mice and the brains were analyzed 1 day later (Figure 1f), tdTomato-positive cells were again detected exclusively in *Ai9/+; Glast*^*CreERT2/+*^ mice, but not in control *Ai9/+* mice (Figure 1g). All tdTomato-positive cells were SOX9-positive astroglial lineage cells (Figure 1h, i). Nearly all tdTomato-positive cells were BG-like cells (BGLPs and BGs; SOX9+VIMENTIN+ unipolar cells located in the PCL and ML; 96.6 ± 0.6%), whereas a small fraction were astrocytes in the IGL or WM (SOX9+VIMENTIN+KI67− multipolar cells located in the IGL or WM; 3.4 ± 0.6%) (Figure 1h, j). As observed at P6, AsLPs (SOX9+VIMENTIN+KI67+ multipolar cells) were not labeled at P0 (Figure 1h, j). Together, these results indicate that tdTomato-positive cells labeled by cerebellar surface application of Tx were restricted to SOX9-positive astroglial lineage cells and were predominantly BG-like cells (BGLPs and BGs). These findings suggest that, among cells labeled by this tamoxifen-based surface-labeling approach, BGLPs are the only proliferative population. These results established that cerebellar surface Tx application predominantly labels BG-like astroglial cells at P0 and P6. We therefore used this strategy to compare the lineage output of BGLPs labeled at multiple postnatal stages (Figures 2 and 3).

**Figure 3.**
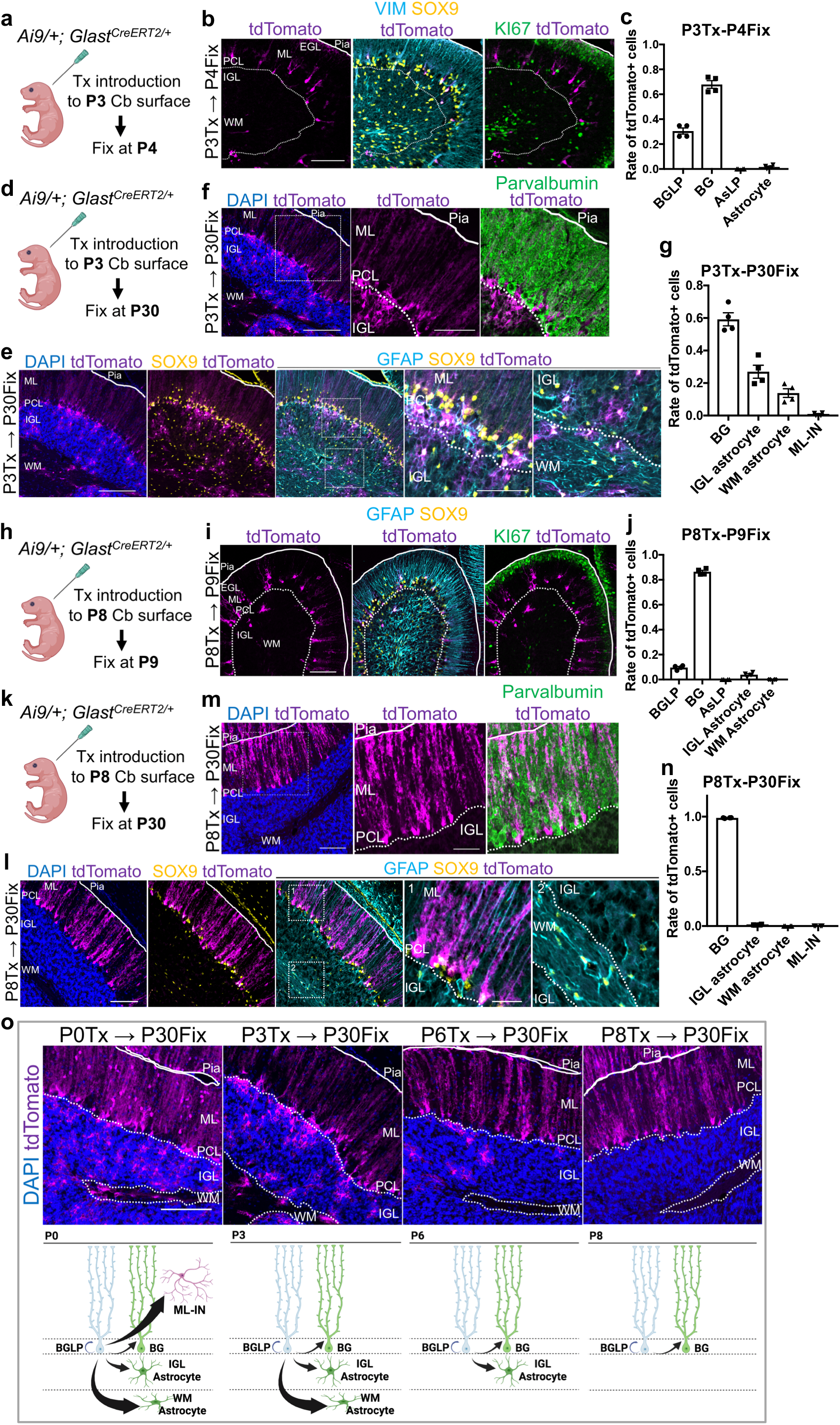
BGLPs undergo progressive lineage restriction during postnatal development. (a) Experimental scheme. Tx were applied to the cerebellar surface of *Ai9/+; Glast*^*CreERT2/+*^ mice at P3, and the mice were fixed at P4. (b) Immunostaining of cerebella from tamoxifen-treated *Ai9/+; Glast*^*CreERT2/+*^ mice for KI67, SOX9, VIMENTIN (VIM) and tdTomato. Scale bars: 50 μm. (c) Quantification of the identities of tdTomato-positive cells. The definitions of astroglial lineage cells (BGLPs, BGs, AsLPs, and astrocytes) were the same as those in Figure 1j. N = 4 mice. (d) Experimental scheme. Tx were applied to the cerebellar surface of *Ai9/+; Glast*^*CreERT2/+*^ mice at P3, and the mice were fixed at P30. (e, f) Immunostaining of cerebella from tamoxifen-treated *Ai9/+; Glast*^*CreERT2/+*^ mice for DAPI, SOX9, and GFAP (e), or DAPI, Parvalbumin (f), and tdTomato. Scale bars: 100 μm for the lower-magnification images in (e) and (f), 50 μm for the higher-magnification images in (e), and 60 μm for the higher-magnification images in (f). (g) Quantification of the identities of tdTomato-positive cells. The definitions of cells were the same as those in Figure 2h. N = 4 mice. (h) Experimental scheme. Tx were applied to the cerebellar surface of *Ai9/+; Glast*^*CreERT2/+*^ mice at P8, and the mice were fixed at P9. (i) Immunostaining of cerebella from tamoxifen-treated *Ai9/+; Glast*^*CreERT2/+*^ mice for KI67, SOX9, and GFAP. Scale bars: 100 μm. (j) Quantification of the identities of tdTomato-positive cells. The definitions of astroglial lineage cells (BGLPs, BGs, AsLPs, IGL astrocytes, WM astrocytes) were the same as those in Figure 1e. N = 4 mice. (k) Experimental scheme. Tx were applied to the cerebellar surface of *Ai9/+; Glast*^*CreERT2/+*^ mice at P8, and the mice were fixed at P30. (l, m) Immunostaining of cerebella from tamoxifen-treated *Ai9/+; Glast*^*CreERT2/+*^ mice for DAPI, SOX9, and GFAP (l), or DAPI, Parvalbumin (m), and tdTomato. Scale bars: 100 μm for the lower-magnification images in (l) and (m), 30 μm for the higher-magnification images in (l), and 50 μm for the higher-magnification images in (m). (n) Quantification of the identities of tdTomato-positive cells. The definitions of astroglial lineage cells and ML-INs were the same as those in Figure 2h. N = 5 mice. All data are shown as mean ± SEM. (o) Model summarizing the progressive lineage restriction of BGLPs during postnatal development. The upper panels show representative images of DAPI and tdTomato staining in mice treated with Tx at the indicated postnatal stages and fixed at P30, and the lower panels show the corresponding schematic models. P0 BGLPs generate BGs, IGL astrocytes, WM astrocytes, and ML-INs. P3 BGLPs generate BGs, IGL astrocytes, and WM astrocytes, but not ML-INs. P6 BGLPs generate BGs and IGL astrocytes, but not WM astrocytes or ML-INs. P8 BGLPs generate predominantly BGs.

### Recombination-based lineage tracing confirms broader lineage output of P0 than P6 BGLPs

Having established the specificity of the surface Tx-labeling approach, we next used recombination-based lineage tracing to test whether P0 BGLPs show broader lineage output than P6 BGLPs (Figure 2a), as suggested by our previous electroporation-based study (Suyama et al., 2026). At both labeling stages, tdTomato-positive cells were detected exclusively in *Ai9/+; Glast*^*CreERT2/+*^ mice, but not in control *Ai9/+* mice (Figure 2b, c). When we analyzed the mice labeled at P6, all tdTomato-positive cells were SOX9-positive; among them, 81.6 ± 0.9% BGs (SOX9+GFAP+ unipolar cells located in the PCL and ML) and 17.6 ± 0.7% IGL astrocytes (SOX9+GFAP+ multipolar cells located in the IGL) (Figure 2d, h). In contrast, WM astrocytes (SOX9+GFAP+ multipolar cells located in the WM), ML-INs (Parvalbumin+ cells located in the ML), Purkinje cells (PCs; CalbindinD28k+ cells), granule cells (GCs; PAX6+ cells), unipolar brush cells (UBCs; TBR2+ cells), and oligodendrocyte lineage cells (OL-lineage cells; OLIG2+ cells) were all tdTomato-negative (Figure 2d–h). These results were consistent with those reported by Cerrato and colleagues using a recombination-based method (Cerrato et al., 2018).

We next analyzed the mice labeled at P0. Among tdTomato-positive cells labeled at P0, 76.4 ± 2.2% were SOX9-positive, consisting of 49.5 ± 2.2% BGs, 16.6 ± 2.2% IGL astrocytes, and 10.3 ± 0.9% WM astrocytes (Figure 2d, h). By contrast, all tdTomato-positive but SOX9-negative cells were ML-INs (Figure 2e–h). PCs, GCs, UBCs, and OL-lineage cells were all tdTomato-negative (Figure 2f–h). Thus, BGLPs labeled at P0 gave rise to ML-INs in addition to astroglial lineage cells. These results were consistent with our previous findings obtained using electroporation-based labeling (Suyama et al., 2026). Together, these results confirmed, using recombination-based lineage tracing, that P0 BGLPs have broader lineage output than P6 BGLPs. The production of ML-INs from P0-labeled BGLPs was further examined in later experiments.

### BGLPs undergo progressive lineage restriction during postnatal development

The P0 and P6 analyses suggested that BGLP lineage output becomes restricted during postnatal development. To determine whether this restriction occurs abruptly or progressively, we next analyzed the progeny of BGLPs labeled at intermediate and late postnatal stages, P3 and P8 (Figure 3). We first applied Tx to the cerebellar surface of P3 *Ai9/+; Glast*^*CreERT2/+*^ mice (Figure 3a–g). When the brains were analyzed 1 day later (Figure 3a), as observed after Tx application at P0 or P6, all tdTomato-positive cells were SOX9-positive astroglial lineage cells (Figure 3b, c). The vast majority of these cells were BG-like cells (BGLPs or BGs) (98.4 ± 0.6%), whereas a small fraction were astrocytes in the IGL or WM (1.6 ± 0.6%) (Figure 3b, c). We next applied Tx at P3 and analyzed the brains at P30 (Figure 3d). All tdTomato-positive cells were SOX9-positive astroglial lineage cells (Figure 3e–g), consisting of BGs (59.2 ± 3.4%), IGL astrocytes (27.0 ± 3.5%), and WM astrocytes (13.8 ± 2.3%) (Figure 3e, g). No ML-INs were detected (Figure 3f, g). These results suggest that P3 BGLPs give rise to BGs, IGL astrocytes, and WM astrocytes, but not ML-INs.

Next, to determine which cell types are generated by BGLPs at P8, a stage immediately before their disappearance, we applied Tx to the cerebellar surface of P8 *Ai9/+; Glast*^*CreERT2/+*^ mice (Figure 3h–n). When the brains were analyzed 1 day later (Figure 3h), as observed at P0, P3, and P6, all tdTomato-positive cells were SOX9-positive astroglial lineage cells (Figure 3i, j). The vast majority of these cells were BG-like cells (BGLPs or BGs) (96.1 ± 0.7%), whereas a small fraction were IGL astrocytes (3.9 ± 0.7%) (Figure 3i, j). We next applied Tx at P8 and analyzed the brains at P30 (Figure 3k). All tdTomato-positive cells were SOX9-positive astroglial lineage cells (Figure 3l–n), and nearly all of them were BGs (98.8 ± 0.1%), whereas a small fraction were IGL astrocytes (1.2 ± 0.1%) (Figure 3l, n). Together with the P0 and P6 analyses shown in Figure 2, these results indicate that BGLP lineage output narrows stepwise during postnatal development: P0, P3, P6, and P8 BGLPs generate four, three, two, and predominantly one progeny category, respectively. This stepwise reduction in lineage output is summarized in Figure 3o.

### ML-INs are likely generated indirectly from P0 BGLPs via AsLPs and INPs

Finally, because P0-labeled BGLPs gave rise to ML-INs, we investigated whether this neurogenic output occurs directly or through intermediate progenitor populations. ML-INs in the mouse cerebellum are generated mainly during postnatal development from inhibitory neuron progenitors (INPs) located in the WM, which are identified by expression of PAX2 or ASCL1 (Joyner & Bayin, 2022; Leto et al., 2006).

By contrast, AsLPs located in the WM are known to be a source of WM astrocytes (Parmigiani et al., 2015), although at least a subset of them has been suggested to interconvert with INPs and contribute to the production of ML-INs (Joyner & Bayin, 2022). Since P0 BGLPs were shown to generate ML-INs, we next examined whether these neurons are generated directly from P0 BGLPs or indirectly via intermediate cell populations such as AsLPs and INPs.

We first applied Tx to the cerebellar surface of P0 *Ai9/+; Glast*^*CreERT2/+*^ mice and analyzed the brains at P3 (Figure 4a). All tdTomato-positive cells were SOX9-positive and consisted of BGLPs (16.9 ± 0.7%), BGs (50.4 ± 2.3%), astrocytes (14.4 ± 1.8%), and AsLPs (17.6 ± 0.8%) (Figure 4b, f). By contrast, neither INPs (PAX2+KI67+ cells) nor differentiated inhibitory neurons (INs; PAX2+KI67− cells) were labeled by tdTomato at this stage (Figure 4c, f). These results indicate that P0 BGLP descendants include AsLPs by P3. Because all tdTomato-positive cells detected at this stage belonged to the astroglial lineage, the earliest detectable descendants of P0-labeled BGLPs appear to be restricted to astroglial lineage cells.

**Figure 4.**
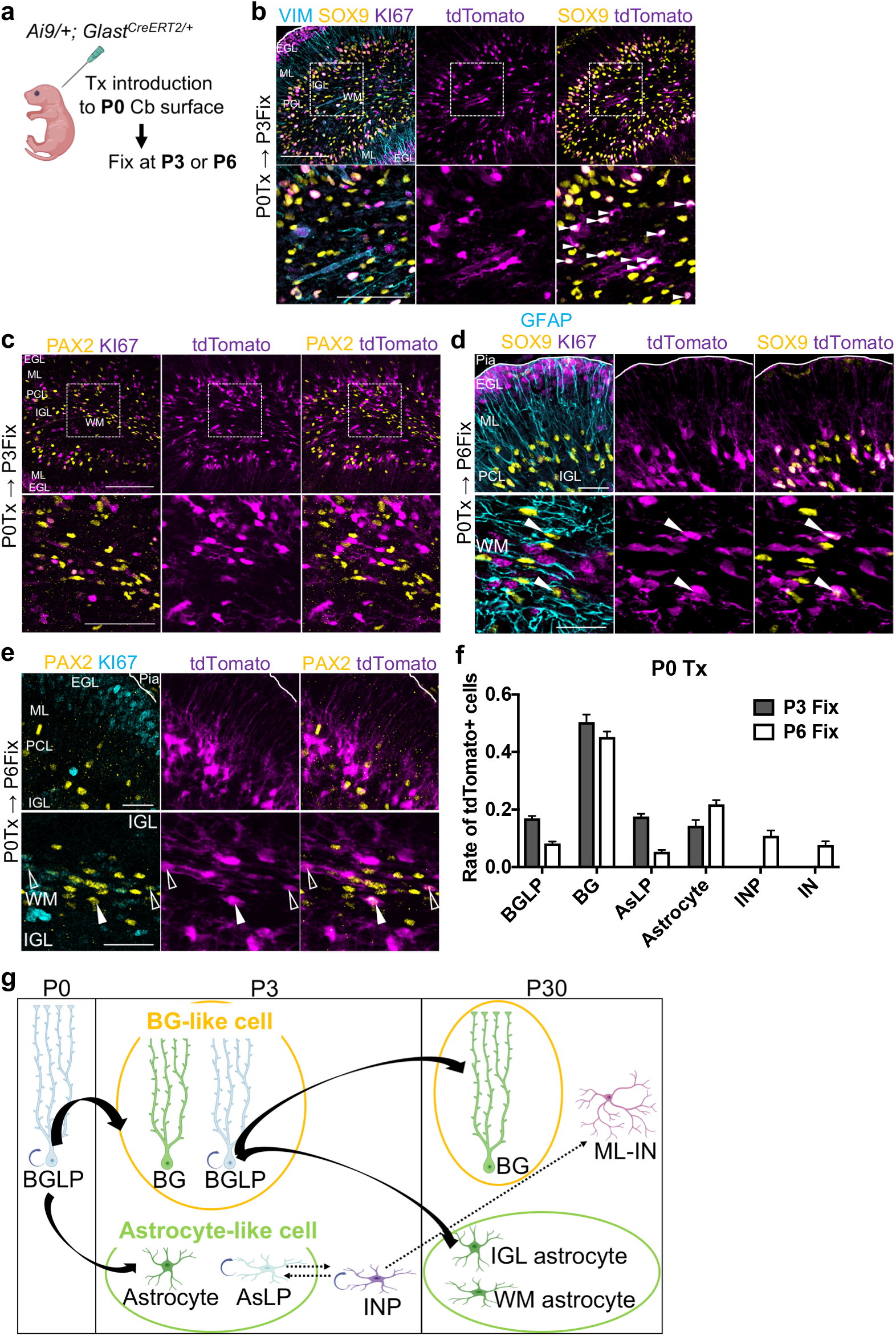
ML-INs are likely generated indirectly from P0 BGLPs via AsLPs and INPs. (a) Experimental scheme. Tx was applied to the cerebellar surface of *Ai9/+; Glast*^*CreERT2/+*^ mice at P0, and the mice were fixed at P3 or P6. (b, c) Immunostaining of cerebella from tamoxifen-treated *Ai9/+; Glast*^*CreERT2/+*^ mice fixed at P3 (P0Tx→P3Fix) for KI67, SOX9, and VIMENTIN (VIM) (b), or KI67, PAX2, and tdTomato (c). Scale bars: 100 μm for the upper panels and 50 μm for the lower panels. White arrowheads in the lower panels of (b) indicate AsLPs or astrocytes (SOX9+VIM+ cells with multipolar processes located in the IGL or WM). (d, e) Immunostaining of cerebella from tamoxifen-treated *Ai9/+; Glast*^*CreERT2/+*^ mice fixed at P6 (P0Tx→P6Fix) for KI67, SOX9, and GFAP (d), or KI67, PAX2, and tdTomato (e). The upper panels show the EGL, ML, PCL, and IGL, whereas the lower panels show the WM. White arrowheads in the lower panels of (d) indicate AsLPs (SOX9+GFAP+KI67+ cells with multipolar processes located in the WM). White arrowheads in the lower panels of (e) indicate INPs (PAX2+KI67+ cells located in the WM), whereas open arrowheads indicate INs (PAX2+KI67− cells located in the WM). Scale bars: 30 μm. (f) Quantification of the identities of tdTomato-positive cells. The definitions of astroglial lineage cells (BGLPs, BGs, AsLPs, and astrocytes) were the same as those in Figure 1j. The definitions of INPs and INs were the same as those in (e). N = 5 mice for the P0Tx→P3Fix analysis and N = 4 mice for the P0Tx→P6Fix analysis. All data are shown as mean ± SEM. (g) Model suggesting that ML-INs are generated indirectly from P0 BGLPs. At P3, P0 BGLP descendants consisted only of BG-like cells (BGLPs and BGs) and astrocyte-like cells (AsLPs and astrocytes) (Figure 4b, c, f). In addition, BGLPs labeled at P3 generated BGs, IGL astrocytes, and WM astrocytes, but not ML-INs (Figure 3d–g, o). By contrast, between P3 and P6, P0 BGLP descendants included not only AsLPs but also INPs and differentiated INs (Figure 4d–f). Because AsLPs are known to interconvert with INPs and to contribute to ML-IN production (Joyner & Bayin, 2022), these observations suggest that ML-INs are generated indirectly from P0 BGLPs via AsLPs and INPs. Solid black arrows indicate relationships supported by the present study, whereas dashed arrows indicate putative lineage relationships inferred from previous studies (Joyner & Bayin, 2022; Leto et al., 2006).

We next applied Tx to the cerebellar surface of P0 *Ai9/+; Glast*^*CreERT2/+*^ mice and analyzed the brains at P6 (Figure 4a). tdTomato-positive cells included BGLPs (8.1 ± 0.5%), BGs (45.2 ± 1.7%), IGL astrocytes (13.4 ± 0.4%), and WM astrocytes (8.4 ± 0.9%), as well as AsLPs (5.3 ± 0.6%) and INPs (10.9 ± 1.6%) (Figure 4d–f). In addition, INs (7.7 ± 1.1%) were also detected (Figure 4e, f). These results indicate that cells derived from P0 BGLPs acquire INP and IN identities between P3 and P6. Together with the finding that P3 BGLPs do not generate neuronal lineage cells (Figure 3e–g), this supports the idea that the cells directly generated from BGLPs are restricted to astroglial lineage cells. By contrast, INPs and INs are likely generated indirectly, possibly via AsLPs, during ML-IN production from P0 BGLPs (Figure 4g).

## Discussion

In this study, we identified postnatal BGLPs as a transient progenitor population that undergoes progressive lineage restriction during cerebellar development. By using recombination-based lineage tracing, we found that the range of cell types generated from BGLPs narrowed stepwise as development proceeded. BGLPs labeled at P0 gave rise to BGs, IGL astrocytes, WM astrocytes, and ML-INs; BGLPs labeled at P3 generated BGs, IGL astrocytes, and WM astrocytes; BGLPs labeled at P6 generated BGs and IGL astrocytes; and BGLPs labeled at P8 generated predominantly BGs. Thus, the lineage output of BGLPs changed from four progeny categories at P0 to a predominantly BG-restricted output by P8. These findings suggest that BGLPs dynamically alter their lineage potential during the short postnatal period in which they are present, thereby contributing to stage-dependent cellular supply in the developing cerebellum.

Our previous electroporation-based study suggested that P0 BGLPs have broader lineage potential than P6 BGLPs, giving rise not only to BGs and IGL astrocytes but also to WM astrocytes and ML-INs (Suyama et al., 2026). However, because electroporation-based labeling is not a permanent recombination-based lineage tracing method, the full progeny of initially labeled BGLPs could not be rigorously defined. In the present study, cerebellar surface tamoxifen application to *Ai9/+; Glast*^*CreERT2/+*^ mice allowed recombination-based labeling of predominantly BG-like cells, among which BGLPs were the only proliferative population. Using this approach, we confirmed the broader lineage potential of P0 BGLPs and the more restricted output of P6 BGLPs. More importantly, by extending the analysis to P3 and P8, we showed that the difference between P0 and P6 BGLPs is not simply a binary stage difference, but part of a continuous process of progressive lineage restriction during postnatal development.

The stepwise narrowing of BGLP lineage potential suggests that BGLPs may provide a useful *in vivo* model for studying how developmental time is translated into progenitor competence. Temporal changes in developmental potential have been described in several neural progenitor populations, including radial glial cells, which generate different neuronal and glial cell types as development proceeds (Koo et al., 2023; Telley et al., 2019). In such systems, progenitor competence is thought to be regulated by changes in transcriptional programs, chromatin states, and extracellular cues (Koo et al., 2023; Telley et al., 2019; Yoon et al., 2018). BGLPs may offer a complementary and experimentally tractable model because they are anatomically identifiable, exist within a short postnatal developmental window, and show a clear reduction in progeny diversity across defined stages. Another practical advantage of this system is that genes can be introduced into BGLPs *in vivo* by cerebellar surface electroporation, enabling relatively simple gain- and loss-of-function analyses during the period in which their lineage potential is changing. In addition, our recent study identified molecular differences between P0 and P6 BGLPs, including higher expression of stem cell-associated and cell cycle-related genes in P0 BGLPs and more BG-like molecular features in P6 BGLPs (Suyama et al., 2026). Together with the present lineage-tracing results, these features make BGLPs a promising model for linking stage-dependent molecular states to progressive lineage restriction. Future studies using this system should clarify which molecular programs drive the transition from broad P0 competence to the predominantly BG-restricted output observed at P8.

Another important observation in this study is that ML-IN production from P0 BGLPs is likely to occur indirectly. When P0-labeled BGLP descendants were analyzed at P3, tdTomato-positive cells included BG-like cells and astrocyte-like cells, including AsLPs, but not INPs or differentiated INs. By contrast, by P6, P0-labeled descendants included not only AsLPs but also INPs and INs. In addition, BGLPs labeled at P3 did not generate ML-INs by P30. These temporal observations support the idea that the earliest detectable descendants of P0 BGLPs are restricted to astroglial lineage cells, and that INPs and INs arise later, possibly through AsLPs. This interpretation is consistent with previous studies suggesting that AsLPs can interconvert with INPs and contribute to ML-IN production. Although the present data do not directly demonstrate the complete lineage sequence from individual BGLPs to AsLPs, INPs, and ML-INs at the single-cell level, they suggest that the glial-neuronal lineage relationship in the early postnatal cerebellum may be more hierarchical and dynamic than previously appreciated.

Several limitations should be considered. First, although cerebellar surface tamoxifen application labeled predominantly BG-like cells, a small fraction of non-BG-like astroglial cells was also labeled. We interpret the long-term progeny primarily as BGLP-derived because BGLPs were the only proliferative population among the initially labeled cells. Nevertheless, future clonal lineage tracing or dual-recombinase approaches would be useful to define the progeny of individual BGLPs more precisely. Second, our classification of BGLPs and BGs relied in part on KI67 expression, and KI67-negative BG-like cells may include both postmitotic BGs and quiescent progenitor-like cells. Third, the proposed indirect pathway from BGLPs to ML-INs through AsLPs and INPs remains inferential. Direct demonstration of this sequence will require approaches that can follow individual intermediate populations over time.

In conclusion, this study demonstrates that BGLPs undergo progressive lineage restriction during postnatal cerebellar development. The range of cell types generated by BGLPs narrows stepwise from broad P0 potential to a predominantly BG-restricted output by P8. This developmental progression suggests that BGLPs contribute to stage-specific cellular supply by dynamically changing their lineage potential over time. Because BGLPs are transient, anatomically identifiable, and exhibit a clear temporal reduction in developmental potential, they may serve as a useful model for investigating the cellular and molecular mechanisms underlying lineage restriction in the developing nervous system.

## Experimental procedures

### Animals

All animal experiments were approved by the Animal Care and Use Committee of the National Institute of Neuroscience, National Center of Neurology and Psychiatry (Tokyo, Japan; Project 202017R2). *Glast*^*CreERT2*^ mice and *Ai9* reporter mice were purchased from The Jackson Laboratory (stock no. 012586 and 007909, respectively) (Mori et al., 2006, Madisen et al., 2010). Mice were maintained under specific pathogen-free conditions on a 12-h light/dark cycle with ad libitum access to food and water. Both male and female mice were used in this study.

### Local administration of tamoxifen crystals

To induce Cre-mediated recombination selectively in BG-like cells (BGLPs and BGs) of *Ai9; Glast*^*CreERT2*^ mice, tamoxifen citrate crystals (Tx; Sigma-Aldrich) were locally applied to the cerebellar surface as previously described, with minor modifications (Cerrato et al., 2018; Parmigiani et al., 2015). Briefly, the skin overlying the posterior head was opened, and a small amount of Tx was placed on the tip of fine forceps. The skull was then gently penetrated with the forceps, and the Tx crystals were directly applied onto the cerebellar surface. Using this procedure, Cre-mediated recombination was induced predominantly in the cerebellum, particularly in BG-like cells (BGLPs and BGs).

### Immunohistochemistry and antibodies

Mice less than 10 days old were decapitated, and their brains were collected and post-fixed in 4% PFA overnight with gentle shaking. Mice older than 10 days were transcardially perfused with fixative, and their brains were collected and post-fixed in 4% PFA overnight with gentle shaking. The brains were subsequently incubated sequentially in 1× PBS, 10% sucrose, and 30% sucrose overnight with gentle shaking. The brains were then embedded in O.C.T. compound (Sakura Finetek). Frozen brains were sagittally sectioned at a thickness of 16 μm using a cryostat (CM3050 S; Leica). Cryosections were washed twice with PBS for 10 min and then incubated at room temperature with 10% normal donkey serum (S30-100ML; Millipore) containing 0.2% Triton X-100 in PBS for 40 min. After blocking, the sections were incubated with primary antibodies diluted in blocking solution at 4 °C for 16 h. The following primary antibodies were used: mouse anti-CalbindinD28K (1:500; C9848; Sigma-Aldrich), rat anti-GFAP (1:500; AB5541; Merck Millipore), chicken anti-GFP (1:1000; ab1218; Abcam), rat anti-Ki67 (1:500; 14-5698-82; Invitrogen), goat anti-NeuroD1 (1:500; AF2746; R&D), goat anti-Olig2 (1:500; AF2418; R&D Systems), rabbit anti-PAX2 (1:500; 71-6000; Invitrogen), mouse anti-Parvalbumin (1:500; 3088; Merck Millipore), rabbit anti-PAX6 (1:500; 901301; BioLegend), goat anti-human SOX9 (1:500; AF3075; R&D Systems), rabbit anti-Tbr2/Eomes (1:1000; ab183991; Abcam), and rabbit anti-Vimentin (1:500; D21H3; Cell Signaling). Subsequently, slides were rinsed with PBS (10 min, twice) and incubated with secondary antibodies conjugated to Alexa Fluor 405, Alexa Fluor 488, Alexa Fluor 568, or Alexa Fluor 647 (1:400; Abcam), or Alexa Fluor 488 (1:400; Jackson), and DAPI (25 μg/mL; Invitrogen) in 0.2% PBST at room temperature for 2 h. The slides were rinsed again with PBS (10 min, twice) and mounted with ProLong Glass Antifade Mountant (P36984; Thermo Fisher Scientific).

### Image acquisition and quantification

The entire sagittal section of the cerebellar vermis was observed under a microscope and analyzed to examine the distribution of tdTomato+ cells in mice at each stage of the experiment. For analysis, all lobules in sagittal sections of the cerebellar vermis were included. For each experimental condition, multiple sagittal sections were analyzed per animal, and at least two images were acquired from each animal. Images were obtained using a confocal microscope, SpinSR10 (Olympus). The acquired images were adjusted and analyzed using ImageJ (RSB, version 2.3.0) or Fiji (RSB, version 1.0) software. All graphs were generated using GraphPad Prism software (GraphPad Software, version 7.0e). Quantification of co-expressed markers in this study was not performed by direct observation under the microscope. Instead, all analyses were conducted using confocal images acquired beforehand under identical imaging conditions. During quantification, each cell was carefully evaluated with reference to DAPI or individual marker staining to ensure correct nuclear assignment, and particular care was taken to avoid counting signals originating from adjacent cells or from different focal planes. This approach minimized potential artifacts arising from signal overlap along the z-axis. To identify astroglial lineage cells, we labeled nuclei with SOX9. Cellular morphology was visualized using antibodies against VIMENTIN (VIM) (at P0-P4) or GFAP (at P6-P30), respectively. KI67 signals were used to discriminate between proliferative progenitors (BGLPs, AsLPs and INPs) and postmitotic cells (BGs, astrocytes and INs). The cerebellum is known to contain minor astroglial populations, including “Fananas-like cells,” which are unipolar astroglial cells located in the molecular layer (ML) (Singer et al., 2025), and “cerebellar nuclei astrocytes,” which are multipolar astroglial cells located in the cerebellar nuclei (Martínez-Mendoza et al., 2023). However, cerebellar surface tamoxifen application did not label these populations in our experiments.

## Acknowledgements

This work was supported by JSPS KAKENHI (Grant Numbers 25K02372, 26H01604 to MH, 21K20853, 23K14203 and 26K18323 to TA); AMED (Grant Numbers 24wm0425005h0004 and 24ek0109764h0001 to MH), an Intramural Research Grant of NCNP (4–5, 4-6, 6-9 to MH); Japan Health Research Promotion Bureau (JH) under Research Fund (2024-D-01 to MH); Multilayered Stress Diseases (JPMXP1323015483 to MH) JST SPRING, (Grant Numbers JPMJSP2180 to KS); Tokumori Yasumoto Memorial Trust (MH) and Takeda Science Foundation (Grants 2024049458 to TA). We thank T.O. and S.M. for fruitful discussions.

## Author contributions

Conceptualization: T.A., and M.H. Methodology: T.A. Investigation: T.A., K.S., S.I., E.I. Formal analysis: T.A. Writing—original draft: T.A. and M.H. Writing—review and editing: All authors. Visualization: T.A. Funding acquisition: T.A., and M.H. Supervision: T.A., M.S., and M.H.

## Data availability

All data supporting the findings of this study are provided within the paper and its Supplementary Information. Any additional information is available from the corresponding author upon reasonable request.

## Code availability

No custom code was generated or used in this study.

## References

Koo B., Lee K.-H., Ming G.L., Yoon K.-J., Song H. Setting the clock of neural progenitor cells during mammalian corticogenesis. Semin. Cell Dev. Biol. 142:43–53. (2023).

Yoon K.-J., Vissers C., Ming G.-L., Song H. Epigenetics and epitranscriptomics in temporal patterning of cortical neural progenitor competence. J. Cell Biol. 217, 1901–1914 (2018).

Telley L., Agirman G., Prados J., Amberg N., Fièvre S., Oberst P., Bartolini G., Vitali I., Cadilhac C., Hippenmeyer S., Nguyen L., Dayer A., Jabaudon D. Temporal patterning of apical progenitors and their daughter neurons in the developing neocortex. Science 364, eaav2522 (2019).

Joyner A.L., Bayin N.S. Cerebellum lineage allocation, morphogenesis and repair: impact of interplay amongst cells. Development 149, dev185587 (2022).

Leto K., Carletti B., Williams I.M., Magrassi L., Rossi F. Different types of cerebellar GABAergic interneurons originate from a common pool of multipotent progenitor cells. J. Neurosci. 26, 11682–11694 (2006).

Li P., Du F., Yuelling L.W., Lin T., Muradimova R.E., Tricarico R., Wang J., Enikolopov G., Bellacosa A., Wechsler-Reya R.J., Yang Z.J. A population of Nestin-expressing progenitors in the cerebellum exhibits increased tumorigenicity. Nat. Neurosci. 16, 1737–1744 (2013).

Suyama K., Adachi T., Mizuno M., Ji K., Isogai E., Hasegawa I., Nishitani K., Sone M., Miyashita S., Owa T., Hoshino M. Molecular characteristics and differentiation control mechanisms of Bergmann Glia-like Progenitors in the Postnatal Mouse Cerebellum. Glia, in press (doi:10.1002/glia.70160).

Cerrato V., Parmigiani E., Figueres-Onate M., Betizeau M., Aprato J., Nanavaty I., Berchialla P., Luzzati F., de’Sperati C., Lopez-Mascaraque L., Buffo A. Multiple origins and modularity in the spatiotemporal emergence of cerebellar astrocyte heterogeneity. PLoS Biol. 16, e2005513 (2018).

Mori T., Tanaka K., Buffo A., Wurst W., Kuhn R., Gotz M. Inducible Gene Deletion in Astroglia and Radial Glia—A Valuable Tool for Functional and Lineage Analysis. Glia 54, 21–34 (2006).

Betizeau M., Cortay V., Patti D., Pfister S., Gautier E., Bellemin-Ménard A., Afanassieff M., Huissoud C., Douglas R.J., Kennedy H., Dehay C. Precursor Diversity and Complexity of Lineage Relationships in the Outer Subventricular Zone of the Primate. Neuron 80, 442–457 (2013).

Madisen L., Zwingman T.A., Sunkin S.M., Oh S.W., Zariwala H.A., Gu H., Ng L.L., Palmiter R.D., Hawrylycz M.J., Jones A.R., Lein E.S., Zeng H. A robust and high-throughput Cre reporting and characterization system for the whole mouse brain. Nat. Neurosci. 13, 133–140 (2010).

Parmigiani E., Leto K., Rolando C., Figueres-Oñate M., Lopez-Mascaraque L., Buffo A., Rossi F. Heterogeneity and bipotency of astroglial-like cerebellar progenitors along the interneuron and glial lineages. J. Neurosci. 35, 7388–7402 (2015)

Singer A., Trigo F., Vinel L., Gruere O., Llano I., Oheim M. A first morphological and electrophysiological characterization of Fañanas cells of the mouse cerebellum. J. Physiol. 603, 855–871 (2025).

Martínez-Mendoza M.L., Rodríguez-Arzate C.A., Gomez-Gonzalez G.B., Rosas-Arellano A., Marinez-Torres A. Morphological characteristics of astrocytes of the fastigial nucleus. Heliyon 9, e18006 (2023).

